# Testing for heterogeneous rates of discrete character evolution on phylogenies

**DOI:** 10.1101/2021.09.14.460362

**Authors:** Liam J. Revell, Klaus P. Schliep, D. Luke Mahler, Travis Ingram

## Abstract

Many hypotheses in the field of phylogenetic comparative biology involve specific changes in the rate or process of trait evolution. We present a method designed to test whether the rate of evolution of a discrete character has changed in one or more clades, lineages, or time periods. This method differs from other related approaches (such as the ‘covarion’ model) in that the ‘regimes’ in which the rate or process is postulated to have changed are specified a priori by the user, rather than inferred from the data. Similarly, it differs from methods designed to model a correlation between two binary traits in that the regimes mapped onto the tree are fixed. We apply our method to investigate the rate of dewlap color and/or caudal vertebra number evolution in Caribbean and mainland clades of the diverse lizard genus *Anolis*. We find little evidence to support any difference between mainland and island evolution in either character. We also examine the statistical properties of the method more generally and show that it has acceptable type I error, parameter estimation, and power. Finally, we discuss the relationship of our method to existing models of heterogeneity in the rate of discrete character evolution on phylogenies.

## Introduction

Over recent decades phylogenetic comparative methods have grown steadily in their importance and now assume a relatively central role in evolutionary research (Harmon 2018). The majority of phylogenetic comparative methods take a phylogenetic tree and phenotypic trait data for the constituent species of that tree with the aim of combining the two to better understand trait evolution over the period of history represented by the phylogeny. These inferences might include, for instance, that two or more phenotypic attributes of a clade tend to evolve in a correlated fashion, or that the phenotypic characteristics of an ancestral species were most likely to have been more similar to one of its extant descendants than to others. In the former case we might use the phylogenetically independent contrasts algorithm of Felsenstein (1985) or Grafen’s (1987) phylogenetic regression; whereas in the latter we could employ Maximum Likelihood or Bayesian ancestral state estimation (e.g., Schluter et al. 1997; Pagel et al. 2004).

Increasingly, we have seen substantial growth in the quantity and diversity of phylogenetic comparative methods designed to explicitly model heterogeneity in the process of evolutionary change among lineages or through time. Some of the simplest of these, such as the ‘early-burst’ model (also known as the ACDC model; Blomberg et al. 2003), allow the evolutionary rate to change as a continuous function of time since the root of the tree. Other more sophisticated approaches (e.g., Rabosky 2014) permit the rate of evolution to vary continuously among lineages, but not necessarily as a function of elapsed time since the global root. Others still model heterogeneity in the evolutionary rate or process via discrete regime-shifts, but in which the probability that any lineage is in any regime is either inferred explicitly from the data or integrated over during the analysis (e.g., Revell et al. 2012; Beaulieu et al. 2013; Mahler et al. 2013; Uyeda and Harmon 2014).

Finally, an important class of approach involves the explicit a priori specification by the user of the different regimes that are hypothesized to evolve heterogeneously (e.g., Butler and King 2004; O’Meara et al. 2006). In this case the idea is fairly simple. The investigator begins by fixing a set of regimes on the tree based on a biological hypothesis for how evolution may have proceeded in their group of interest. These regimes could correspond to clades, time periods, individual branches of the phylogeny, or to the postulated history of a discrete character mapped onto the nodes and branches of the phylogeny. One then proceeds to fit a model in which the process of evolution, or the parameter values that describe evolution via this process, is permitted to differ between regimes.

For a number of years this type of approach has been quite popular among comparative biologists. Surprisingly, however, this specific class of method has mostly been applied, to date, for continuously-valued character traits (but see Yang 1998; Yang and Nielsen 1998; below). For instance, in 2004 Butler and King presented a method in which different clades of the tree were permitted to evolve via an Ornstein-Uhlenbeck process with different values of the parameter *θ*, often biologically interpreted as the optimal value of the trait under a model chosen to approximate adaptive evolution (Butler and King 2004). Slightly later, O’Meara and colleagues (also see Thomas et al. 2006) presented a related method in which the rate of evolution via Brownian motion was permitted to vary among branches and clades of the tree according to a hypothesis specified a priori by the user (O’Meara et al. 2006). Revell and Collar (2009) slightly extended this approach to multiple potentially covarying characters. Finally, Beaulieu et al. (2012) presented a highly flexible framework for modelling heterogeneous continuous trait evolution on the tree in which all parameters of an Ornstein-Uhlenbeck or Brownian process are permitted to vary with any configuration of an arbitrary number of hypothesized regimes mapped onto the phylogeny.

Although discretely-valued character traits are often studied in phylogenetic comparative research, to our surprise no precisely comparable set of methods had yet been developed for discrete characters. Perhaps the most similar model is one denominated the codon ‘branch model’ by Yang (1998; Yang and Nielsen 1998, 2002). According to this model, different pre-specified edges of an independently estimated phylogeny are permitted to evolve with different nonsynonymous/synonymous (*d*_N_/*d*_S_) nucleotide substitution rate ratios. This scenario could then be compared to a simpler model in which *d*_N_/*d*_S_ ratio does not vary among the edges and nodes of the phylogeny. To our knowledge, however, this model has never been extended beyond its original intent as a test for positive selection on gene sequences (Yang 1998).

That being said, several related approaches have been proposed for discrete phenotypic traits and are in wide use. For instance, Pagel (1994) presented a method in which the rate of one binary trait is permitted to vary as a function of the state of a second (and, possibly, vice versa). Since this model will fit well when two different discrete characters have a disproportionate tendency to evolve towards certain trait combinations, the model is often interpreted as one that can be used to measure the correlated evolution of discrete characters (but see Maddison and Fitzjohn 2015). Later, Tuffley and Steel (1997) descrixbed a model for molecular evolution which they dubbed the ‘covarion model.’ According to the covarion model a discrete character evolves via a process with two hidden (that is, unobserved and unknown a priori) rate categories: either ‘on’ (in which it can change state) or ‘off’ (in which it cannot; Tuffley and Steel 1997; Penny et al. 2001). This was extended by Galtier (2001) to permit any number of hidden rate categories, and subsequently generalized and applied to phylogenetic comparative analysis by Beaulieu et al. (2013). All of these latter methods suppose that the rate or process of one discrete trait varies among the edges and nodes of our phylogeny either as a function of the observed (e.g., Pagel 1994) or unobserved state of second discretely-valued character (e.g., Penny et al. 2001; Beaulieu et al. 2013).

Herein, we imagine a scenario in which the process of evolution for a discrete trait varies among a series of a priori postulated regimes. These could be clades of the phylogeny, geological or historical time periods, specific edges in which an increase or decrease in rate is hypothesized, or the known (rather than reconstructed) history of a second discrete character trait. Note that this approach should not be taken as an alternative to the aforementioned methods which either seek to identify heterogeneity in the evolutionary process for one character due to an unobserved factor or trait; or integrate over uncertainty in the history of an observed discrete character that affects the rate or evolutionary process of a second. Instead, the method of this study should be employed to investigate scenarios in which a particular set of regimes are hypothesized a priori (on biologically justifiable grounds), at which point our model can be fit and compared to, for instance, a simpler scenario of homogeneous evolutionary rates through time, or perhaps to an alternative hypothesis for rate heterogeneity on the phylogeny.

After we describe our new approach to analyzing the evolution of a discretely valued character on the tree, we will proceed to present some relatively simple simulations examining its general statistical performance in terms of type I error, parameter estimation, and power. We will then use the method to test specific, a priori hypotheses about character evolution for two different discrete character traits in the neotropical lizard genus, *Anolis*: dominant dewlap color; and total number of caudal (tail) vertebrae.

## Model, Methods, and Results

### The model

Like nearly all modern methods for studying the evolution of discretely-valued character states on the phylogeny (but see Felsenstein 2005, 2012; Revell 2014) the model of this study is a flavor of the M*k* model of Lewis (2001). The M*k* model is so called because it describes a continuous-time discrete-state Markov chain with a total of *k* possible states. (Thus an M*k* model with two states is sometimes called an M2 model; a model with three states an M3 model; and so on.) Under this model a set of non-negative real numbers (*q*_*i,j*_) gives the instantaneous transition rates between states *i* and *j* for all *i* ≠ *j*.

In the simplest case, we could imagine an M2 process in which *q*_0,1_ = *q*_1,0_: that is to say, the rates of transition from state 0 to 1 and from state 1 to 0 are equal. In this scenario, the probability of beginning in state 0 and ending in state 1 after time *t* can be written as:

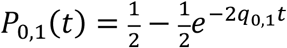

(Lewis 2001). This probability integrates over all the ways in which a time period of length *t* can begin in state 0 and change to state 1: that is, by changing state once, 0 → 1; by changing state, reversing back, and then changing again (0 → 1 → 0 → 1); and so on (Lewis 2001). For conditions in which the probability of multiple changes during time *t* is very small (e.g., low *q*_0,1_, small *t*, or *q*_1,0_ = 0), this expression will converge on the simple integral of an exponential distribution with shape parameter *q*_0,1_ from 0 → *t*, as in this case we are merely computing the probability that a rare event has occurred after time *t* which should have an exponential probability density if the rare event occurs randomly at a constant rate.

Obviously, the probability of starting and ending in the same state 0 is merely one minus our previous expression, or:

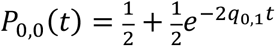

(Lewis 2001). More generally, in a *k*-state process in which transition rates need not be equal nor symmetrical among states, the matrix of transition probabilities between states can be obtained by exponentiating a transition matrix, **Q**, multiplied by the elapsed time, *t*, in which each *i,j*th element of **Q** (for *i* ≠ *j*) contains the instantaneous transition rate between *i* and *j*, and in which the diagonal elements are set to 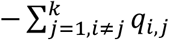, such that each row-sum of **Q** is equal to zero. In other words:

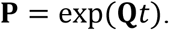

Here, exp(*x*) denotes the matrix exponential of *x* and **P** is a matrix containing the probabilities of changing (or not, on the diagonal of the matrix) between any pair of states. It is straightforward to then proceed and compute the likelihood of any particular value of **Q**, given the data at the tips of the phylogeny and our set of edge lengths, which can be done efficiently using the pruning algorithm of Felsenstein (1973, 1981). For any particular tree and vector of observations for a discrete character at the tips of the tree, the value of **Q** that maximizes this likelihood would be our Maximum Likelihood Estimate (MLE) of the transition matrix **Q**.

(Note that the model called the M*k* model by Lewis 2001 explicitly assumes that all transitions between states occur at the same rate. Although the more general model in which transitions between different pairs of states are permitted to occur at different rates is now often also called the ‘M*k* model’ – in some places it has been more precisely described as the ‘extended’ M*k* model; e.g., Harmon 2018. Herein, we will follow the more typical contemporary convention and refer to the general, *k*-state Markov model as the M*k* model.)

Elaborating on this model but slightly, we imagine two different transition processes, *a* and *b*, operating simultaneously in different parts of the tree in which (for simplicity) 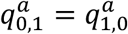 and 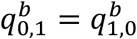, but 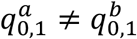. In this case, for a time interval *t* consisting first of time in condition *a, t*_*a*_, followed by time in condition *b, t*_*b*_, the probability of starting in state 0 and ending in state 1 after time *t* = *t*_*a*_ + *t*_*b*_ will be equal to the probability of starting in state 0 and ending time *t*_*a*_ in state 1, then beginning and ending time *t*_*b*_ in state 1, plus the probability of starting and ending time *t*_*a*_ in state 0, plus the probability of starting time interval *t*_*b*_ in state 0 and ending it in state 1. Simply following our equations from earlier, this can be written as:

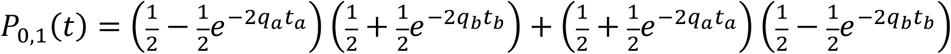

in which I have substituted *q*_*a*_ for 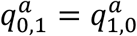 and *q*_*b*_ for 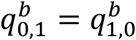. This calculation is illustrated for a simplified ‘phylogeny’ with two taxa in Figure 1, in which the likelihood that is computed (at the root of the tree) is equal to probability of the observed data at the tips (taxon *A* in state ‘0’ and taxon *B* in state ‘1’), given the phylogeny and the model of evolution in which a regime shift (from transition rate *q*_*a*_ to *q*_*b*_) has been postulated along the edge leading to taxon *A*.

**Figure 1.**
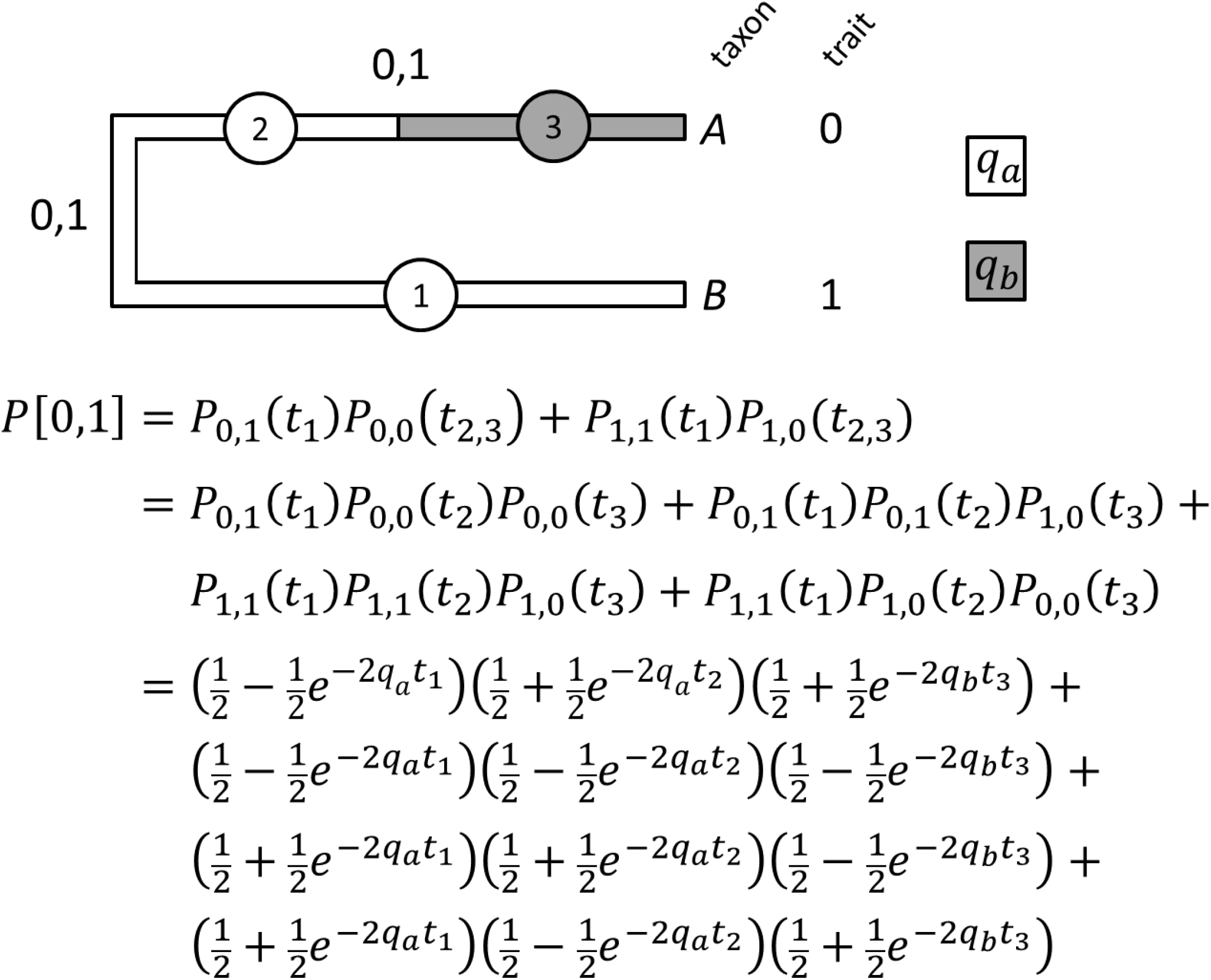
Illustration of the calculation of the probability of observing the data pattern [0,1] for taxa *A* and *B*, respectively, on a two taxon tree with a rate shift hypothesized a priori along one of the two edges of the tree. This is equivalent to the likelihood at the root of the phylogeny. For purposes of simplifying the calculation, we assume that for a given regime (*a* or *b*) the rate of transition (*q*) between binary states 0 and 1 is the same as between states 1 and 0 (that is to say, that 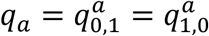 and 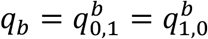); and that states 0 and 1 are equiprobable at the root.

More generally, for arbitrarily different transition matrices **Q**_*a*_ and **Q**_*b*_, the matrix of transition probabilities along a given edge divided into time intervals *t*_*a*_ and *t*_*b*_ can be written as:

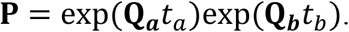

To see how this might be so, let’s once again revert back to our binary character in which we said:

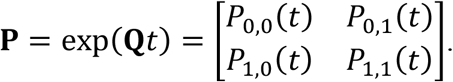

Thus for two successive time periods (*t*_*a*_ and *t*_*b*_) on an edge we have:

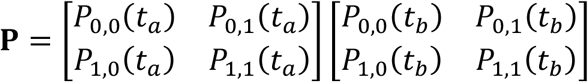

and consequently:

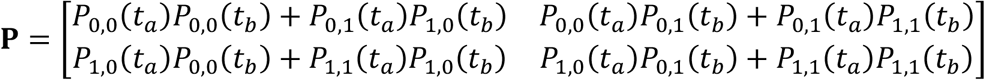

(via normal matrix multiplication), which clearly enumerates all the ways of getting between the two character states over the time intervals *t*_*a*_ and *t*_*b*_. For instance, *P*_1,2_ (the element of **P** in the 1^st^ row and 2^nd^ column) contains the probability of starting *t* in state 0 and ending in state 1, which is equal to the sum of the probability of starting and ending segment *a* of length *t*_*a*_ in state 0, but then changing from 0 to 1 along segment *b* of length *t*_*b*_, plus the probability of starting segment *a* in state 0 and ending state 1, and then not changing state along segment *b*. Note that the calculation of the probability *P*(*i*|*i*) integrates over the possibility of not changing at all and the possibility of one or more changes to other states, followed by a reversal to *i*. Naturally, this can be generalized to any number of character states or rate regimes.

As before, it is straightforward to compute the likelihood for any set of values in **Q**_*a*_ and **Q**_*b*_, using the pruning algorithm of Felsenstein (1973, 1981), and if we identify the values of **Q**_*a*_ and **Q**_*b*_ (and so on, for more than two regimes) that maximize the likelihood, we have found our MLEs of the two or various transition matrices of our model.

### Simulation tests of the method and results

To examine the type I error rate of the method, we first simulated sets of 200 pure-birth phylogenies, each containing either *N* = 25, 50, 100, 200, 400, or 800 taxa, all of which were rescaled to have a total depth of 1.0 unit. We then randomly selected two non-nested clades of each tree with the intent of assigning the tips and edges of these two clades into the same second derived regime, to be then compared against the basal regime on the rest of three. We thus selected the two clades such that: (1) each selected clade contained at least two taxa; and (2) collectively the two clades comprised neither less than 25% nor more than 75% of the tips of the tree. We simulated a constant rate of character evolution on the tree in which *q*_0,1_ = *q*_1,0_ = 0.5, corresponding to an expected number of changes from the root to any tip of 0.5, regardless of the number of taxa in the tree. (Obviously, larger trees nonetheless possessed more changes on average, because they have more total edge length for a given depth.) We then proceeded to fit the heterogeneous rate model presented in this paper along with a standard M*k* (i.e., M2) model with symmetrical transition rates between states. We compared the likelihoods of the two models using a likelihood-ratio test with one degree of freedom, for the one additional parameter estimated in the multi-regime model. The distribution of P-values for each simulation condition is given in supplementary appendix Figure S1; whereas supplementary appendix Table S1 shows the measured type I error rates for each size of tree. On average, we found that type I error was slightly below its nominal level (average type I error = 0.047). Furthermore, in no simulation conditions was type I error significantly greater than 0.05 according to a one-tailed binomial test, and there was no particular tendency for type I error to be higher in smaller or larger phylogenies (Figure S1; Table S1).

We also undertook a set of simulations to examine the power and parameter estimation of the method. For this analysis, we used only the set of 200 simulated 100-taxon trees from the type I error analysis. On these trees we simulated two different rate regimes for the binary character, in which the regime was determined by our a priori mapping and where 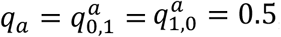, whereas 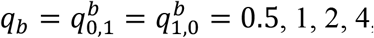, and 8. As one might expect, power to reject the null increased with the difference in rate between regimes (Table S2). The median estimated value of the transition rates was unbiased across all differences in rate (Table S2; Figure 2); however, the mean estimated value of *q*_*b*_ was upwardly biased for higher generating values of *q*_*b*_, evidently due to a small fraction of simulations in which *q*_*b*_ was badly overestimated (Figure 2). We believe that this occurs when by chance there are so many character changes under the faster evolving regime *b* that the data at the tips of any subclade of the tree in state *b* are nearly random with respect to the phylogeny – a result that is only expected if the rate of evolution is very high.

**Figure 2.**
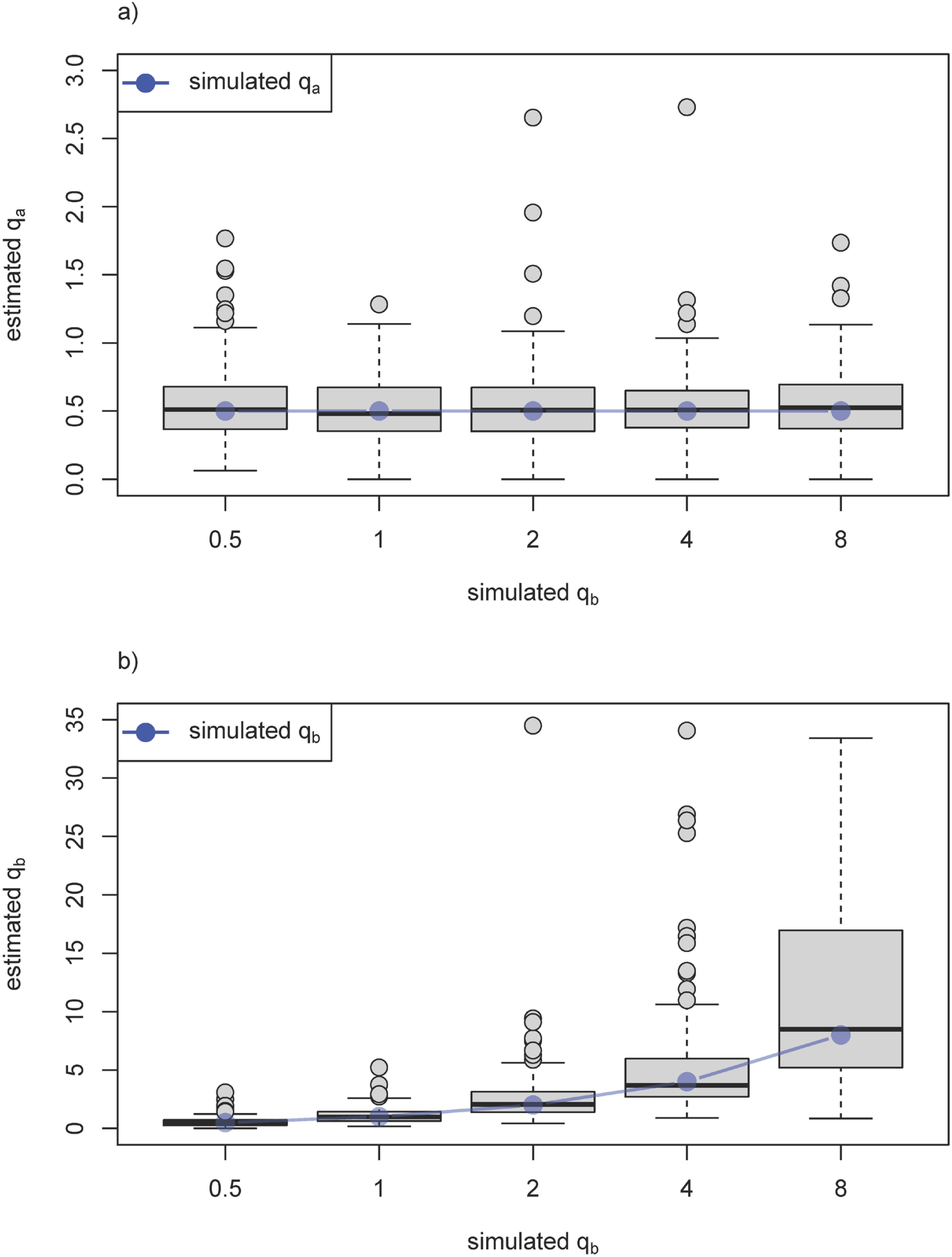
Results from an analysis of parameter estimation. Each panel shows a boxplot for estimated values of the transition rate between states under regime *a* (panel a) and under regime *b* (panel b). As in a standard boxplot, the horizontal black lines show the median value of 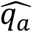 or 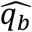 for each simulation, the grey box shows the 25-75% interquartile range, and the whiskers extend 1.5 times the interquartile range or to the maximum or minimum value of 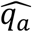 or 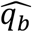 (whichever is smaller or larger, respectively). Values outside 1.5 times the interquartile range are plotted as points. In panel b), 3.7% of points (74 in 2000 simulations) had estimates, 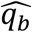, greater than the vertical height of *y* and were thus left out of the plot; however, these values were included in the calculation of the median and interquartile ranges of the boxplot, and in the mean and medians presented in supplementary appendix Table S2. The blue points show the simulated values of *q*_*a*_ and *q*_*b*_ for panels a) and b), respectively.

### Empirical example

In addition to these simulations, we also fit our model to two different empirical cases. In both of these, we examined the rate of evolution of a discrete character in mainland vs. island lineages of lizards in the genus *Anolis*, known as anoles. Since the number of transitions from mainland to island in anoles (and vice versa) is relatively few, we treated these for our purposes as having occurred at known locations on our phylogeny, which we obtained from Gamble et al. (2014). In particular, we assumed that the global ancestral node of all anoles was present on the continental mainland, that occupancy of the Caribbean islands from mainland lineages (or vice versa) occurred via colonization, and then we proceeded to place colonization events precisely halfway along the edge leading to each clade in which descendants were present in the islands. We also reconstructed one island to mainland colonization event, and within this clade a further secondary colonization of islands. (Though this lattermost colonization is not recovered in several other phylogenetic analyses of anoles, e.g., Alföldi et al. 2011, Poe et al. 2017, we do not expect that this detail would affect our analyses.) The mainland/island history that we assumed for the purposes of this analysis is mapped onto the tree of Figure 3.

**Figure 3.**
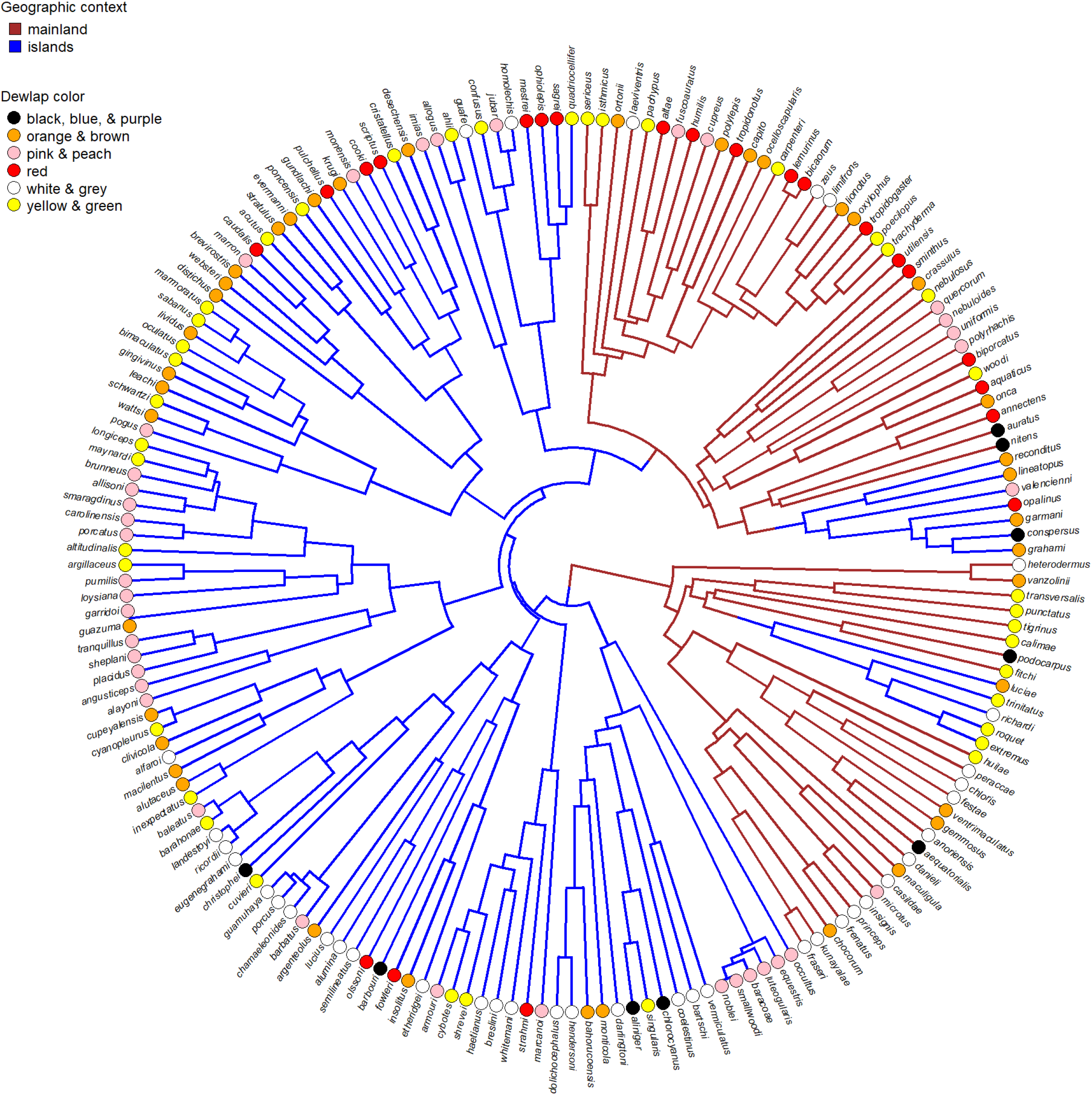
Dominant dewlap color mapped onto a phylogeny of Caribbean (blue branches) and mainland (brown branches) *Anolis* lizards.

Using this mainland/island history as basis for all subsequent inferences, we next analyzed dewlap color evolution. Data for this analysis were originally obtained for a study on dewlap size evolution in anoles (Ingram et al. 2016). The dewlap is an extensible gular fan used by anoles (and some other lizards) for both intra- and interspecific displays. Importantly, the color of the dewlap has been shown to be a critical mate recognition cue in many species (Losos 2009). Our logic in comparing the rate of anole dewlap color evolution between island and continental faunas was the following. Although little studied, at a coarse scale mainland and island anole faunas differ in that mainland communities support higher syntopic species richness but lower average abundances compared to island communities (Andrews 1979; Anderson and Poe *In press*). The ultimate causes of dewlap color diversity remain poorly known (Nicholson et al. 2007; Losos 2009), but if either sexual selection at high densities or selection for species recognition at high diversities is responsible for phylogenetic transitions in dewlap color, we might expect mainland and island anole lineages to exhibit different rates of dewlap color evolution. Dewlaps come in many colors and color combinations, but to keep the discrete character models tractable, we coded dewlap color by placing the dominant (by dewlap surface area) color of each species’ dewlap as being subjectively closest to one of the following six states: *black* (anoles with black, blue, or purple dewlaps); *orange* (orange or brown dewlaps); *pink* (pink and peach dewlapped anoles); *red*; *white* (white and grey dewlaps); or, finally, *yellow* (yellow and green dewlaps; Figure 3). Obviously, considerable nuance will be lost first in reducing the often complex dewlap coloration to a single dominant color for each species, and further still by reducing the genuine interspecific variation in dominant color to our minimal, six-state set. We would not, however, expect this simplification of our dataset to lead to elevated type I error of the method. (To the contrary, we suspect that the opposite is more likely to be true.)

We then proceeded to fit a series of six models to the data and tree. These six models consisted of an evolutionary process in which: transitions occurred at the same rate between all pairs of states (ER); transitions occurred at the same backward and forward rate between each pair of states, but could occur at different rates between different state pairs (SYM); and transitions occurred at different rates between each pair of states (ARD). We fit each of these models either allowing for different rates between mainland and island lineages (-M) or forcing them to have the same rates of change between character states (-S), thus resulting in six models of varying complexity in all (ER-S, ER-M, SYM-S, SYM-M, ARD-S, and ARD-M). Results from this analysis are given in Table 1. In general, although in the best-fitting multi-rate models (SYM-M) the average transition rate between states was higher on islands than in mainland lineages – penalizing for the number of parameters to be estimated, the best-supported model was clearly a model in which both mainland and island fauna dewlap dominant color evolved under the same set of rates of transition between states (SYM-S; Table 1).

**Table 1.**
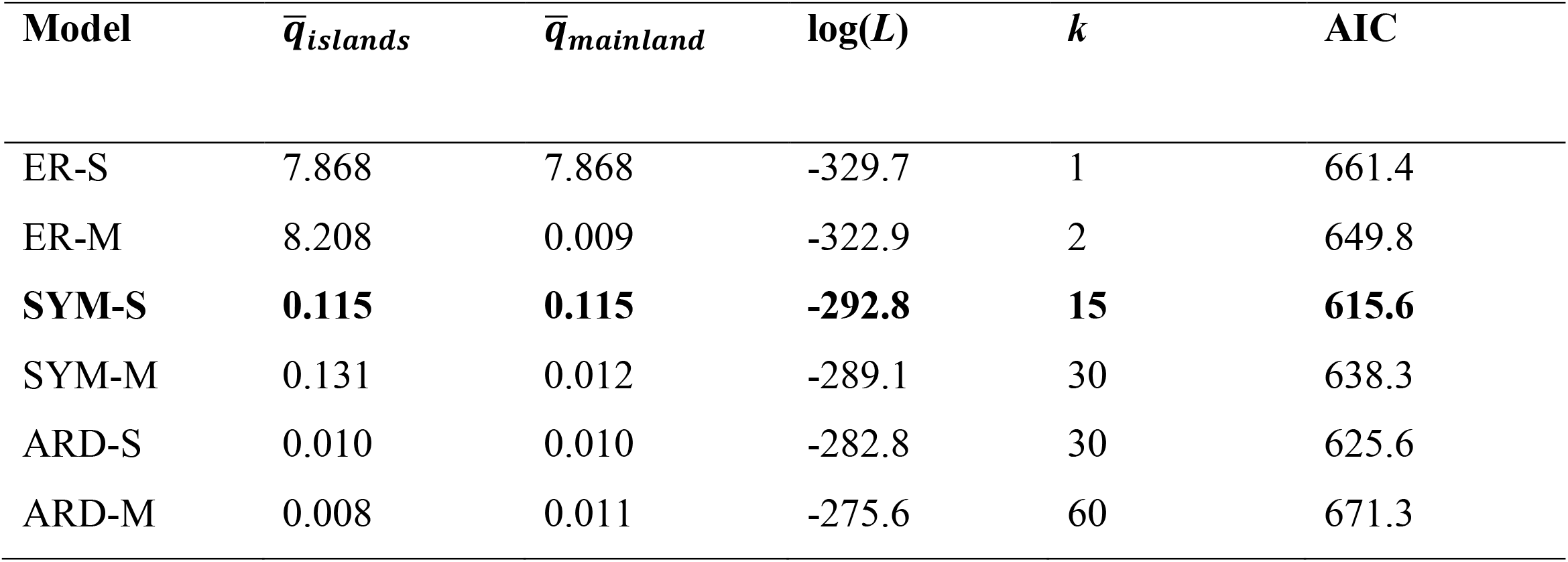
Mean transition rates, log-likelihoods, number of fitted parameters, and AIC for the six fitted models of dewlap dominant color evolution described in the main text. The best supported model (SYM-S) is highlighted in bold text.

| Model | $\bar{q}_{islands}$ | $\bar{q}_{mainland}$ | $\log(L)$ | $k$ | AIC |
| --- | --- | --- | --- | --- | --- |
| ER-S | 7.868 | 7.868 | -329.7 | 1 | 661.4 |
| ER-M | 8.208 | 0.009 | -322.9 | 2 | 649.8 |
| **SYM-S** | **0.115** | **0.115** | **-292.8** | **15** | **615.6** |
| SYM-M | 0.131 | 0.012 | -289.1 | 30 | 638.3 |
| ARD-S | 0.010 | 0.010 | -282.8 | 30 | 625.6 |
| ARD-M | 0.008 | 0.011 | -275.6 | 60 | 671.3 |

In addition to this character, we also analyzed island and mainland caudal (i.e., tail) vertebra number evolution. These data were obtained by simply counting the number of vertebrae from the pelvic girdle to the tip of the tail in a specimen in which the tail was previously deemed to be completely intact (e.g., Figure 4). Our logic in comparing the rate of anole caudal vertebra evolution between island and mainland lineages is simply that conventional wisdom suggests that Caribbean anoles exhibit greater arboreal microhabitat diversification and specialization than do their mainland congeners (Losos 2009; whilst acknowledging that mainland anoles are also ecologically and morphologically diverse, e.g., Pinto et al. 2008). The tail is an appendage that can play an important role in locomotion, particularly in an arboreal setting. Consequently, it seemed reasonable to imagine that it might be under stronger divergent selection in the Caribbean, where anoles may fill a wider array of arboreal and semi-arboreal ecological roles than in continental clades.

**Figure 4.**
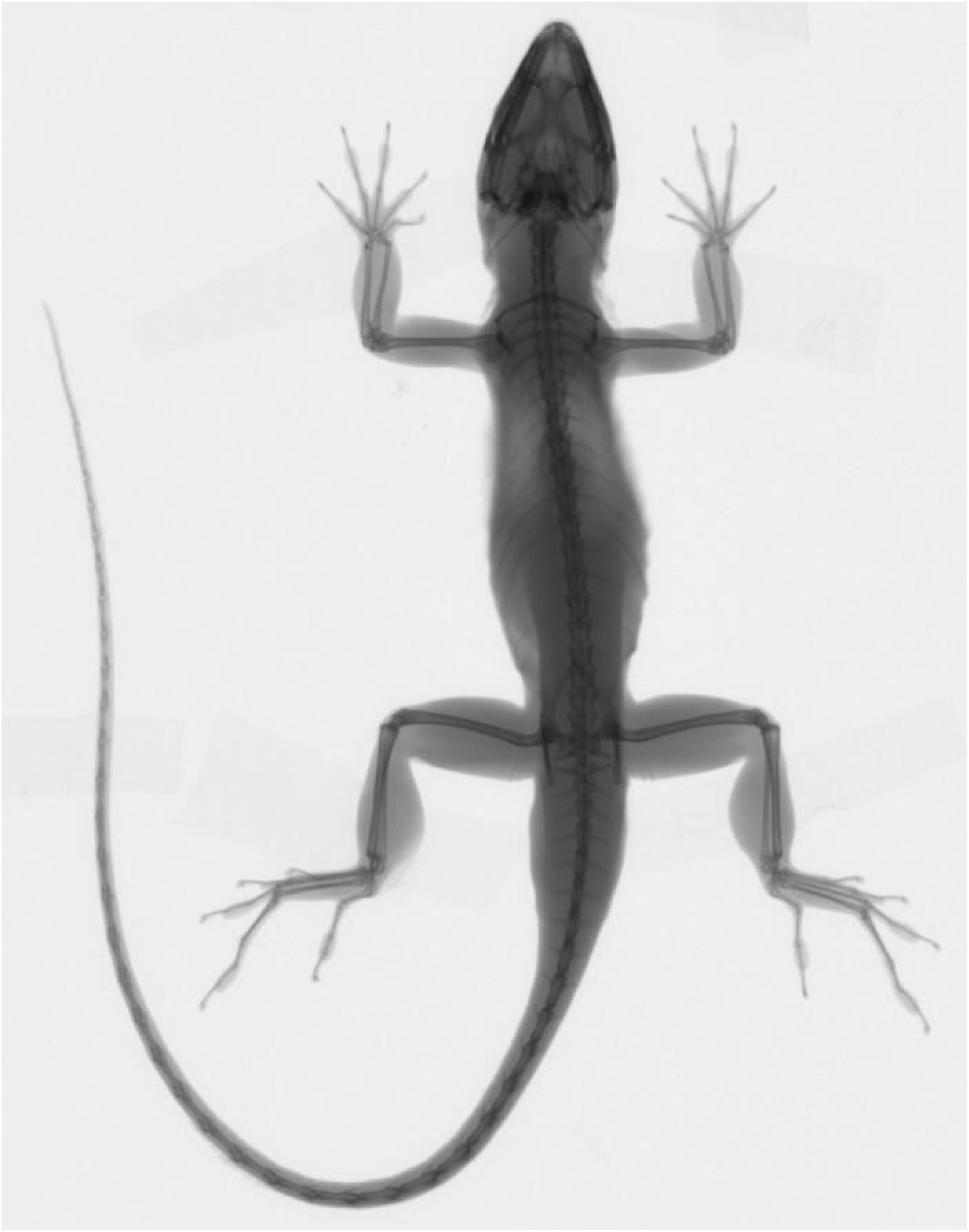
Digital x-ray image of an anole (*Anolis sagrei*) showing the caudal vertebrae of an intact tail. Image courtesy K. Winchell.

Given that the number of caudal vertebrae varies over quite a broad range (from 34 through 55 in these data), one might intuitively assume that the number of parameters to estimate in this model would be impossibly large. In fact, if we make some relatively reasonable simplifying assumptions (keeping in mind that all models are, by definition, intended to be simplifications of reality) the dimensionality of the problem can be quite manageable, even though the state-space is big. Specifically, we decided to treat the acquisition and loss of caudal vertebrae as an ordered process – in which gain and loss were free to proceed with different tempos, but in which changes in the same direction between any pair of adjacent states should occur with the same rate (supplementary appendix Figure S2). Once again, though we found that the estimated rate of character evolution in the best-fitting model was higher in island than in mainland anole lineages, the best-supported model (accounting for parameterization) was, as before, the ordered, single-rate model (Table 2).

**Table 2.** Rate of caudal vertebra loss & gain on the mainland [M] and islands [I], log-likelihoods, number of fitted parameters, and AIC for the two fitted models caudal vertebra number evolution described in the main text. As in Table 1, the best-supported model (ordered-single) is highlighted in bold text.

| Model | $q_{loss}$ | $q_{gain}$ | $\log(L)$ | $k$ | AIC |
| --- | --- | --- | --- | --- | --- |
| **ordered-single** | **0.218** | **0.130** | **-303.4** | **2** | **610.9** |
| ordered-multiple | 0.224 [I] | 0.133 [I] | -303.1 | 4 | 614.3 |
|  | 0.187 [M] | 0.115 [M] |  |  |  |

### Notes on implementation

All the models and methods of this study have been implemented for the R statistical computing environment (R Core Team 2018), and all simulations and analyses were conducted in R. The statistical method described herein has been implemented in the function *fitmultiMk* of the *phytools* R package (Revell, 2012). It is worth noting that this function uses (with attribution and under the Gnu General Public License GPL-3) code to implement the pruning algorithm of Felsenstein (1973, 1981) that was originally published in the *ape* R package of Paradis et al. (2004). *phytools* itself in turn also depends on the important R phylogenetics package *ape* (Paradis et al. 2004; Paradis and Schliep *In press*), as well as on a number of other R packages (Venables and Ripley 2002; Ligges and Mächler 2003; Lemon 2006; Plummer et al. 2006; Harmon et al. 2008; Jackson 2011; Schliep 2011; Chasalow 2012; Xie 2013; Neuwirth 2014; Qiu and Joe 2015; Azzalini and Genz 2016; Becker et al. 2016; Gilbert and Varadhan 2016; Pinheiro et al. 2017). Finally we used the package *lmtest* in some of the analyses and simulation (Zeleis and Hothorn 2002).

## Discussion

In recent decades phylogenetic comparative methods have exploded in popularity within evolutionary research. This growth has generally been accompanied by an appreciation that models assuming a homogeneous process of evolution may be overly simplistic for many phylogenies, particularly those that contain many taxa or that span vast periods of evolutionary time. A number of methods have been developed to explicitly model heterogeneity in the rate or process of phenotypic evolution on the tree. An important class of such method involves the investigator ‘painting’ different a priori regimes onto the branches or clades of a phylogeny, and then testing the hypothesis that evolutionary change for one or more characters differs between different painted regimes. This approach was perhaps best formalized by O’Meara et al. (2006) in the context of testing hypotheses about changes in the rate of evolution for a continuous character evolving by Brownian motion on the tree.

Surprisingly, and in spite of the considerable popularity of this type of analysis for continuous character data, no precisely analogous approach had ever been proposed for discretely valued traits. Herein, we present just such an approach. According to the method of this article, and just as in the method of O’Meara et al. (2006), the user must propose a priori a specific hypothesis for where the rate or process of character evolution is thought to vary on their phylogeny. This hypothesis can then be fit using likelihood, and compared to other hypotheses such as a null hypothesis of constant rates of evolution, or other alternative hypotheses about how the rate varies among clades or branches of the phylogeny. We show that the method, which has been implemented in the R package *phytools* (Revell 2012), has acceptable type I error when the null is true, as well as reasonable power and parameter estimation when it is not (Figures S1, 2; Tables S1, S2).

### Relationship to other methods

As previously noted, models in which regime shifts are postulated a priori by the user, painted on the tree, and then tested against alternative painting or a null hypothesis of homogeneous evolution, have been around for over a decade, and are already quite popular for the analysis of one or more continuous characters on the tree (e.g., Butler and King 2004; O’Meara et al. 2006; Revell and Collar 2009; Beaulieu et al. 2012). We were somewhat surprised to discover that a precise analog did not exist for discretely valued character traits and have tried to fill that void with this contribution. To our knowledge, the sole exception is the codon branch model of Yang (1998; Yang and Nielsen 1998, 2002), which, so far as we are aware, has never been extended beyond testing for positive selection in gene sequences.

Furthermore, as we have attempted to make clear throughout our article and as we reiterate here, in addition to the branch model of Yang (1998) a number of other prior methods have been proposed to explicitly model heterogeneity in the rate or process of evolution in discrete characters on phylogenetic trees. For instance, and perhaps most significantly, Pagel (1994) presented a model in which the rate of evolution of one binary character depends on the state of a second character, or vice versa; or in which the evolution of both characters is interdependent. Since this model will fit well when two characters tend to evolve towards certain character state combinations, the method of Pagel (1994) is often interpreted as a test for the correlated evolution of two binary traits. As shown by Maddison and Fitzjohn (2015; and as is obvious from the structure of the model), Pagel’s (1994) method will also lead to a significant result if a change in value for one character is correlated with a change in the rate of evolution for a second (even if the former is unreplicated on the tree; Maddison and Fitzjohn 2015). Our method too will be significant if our discrete character changes in its rate or process of evolution within a single clade that we have specified a priori. However, in our case this is by design, not by accident. Under this circumstance, a significant result would merely indicate that the data suggest that our single focal subclade evolved by a different process or rate than did the taxa of the rest of the phylogeny, and any attribution of this finding to a specific biological cause is not implied by the result and would be left instead to the interpretation of the investigator.

Another very important class of analytical method also exists for modeling rate heterogeneity of a discrete character trait on the phylogeny. These are models in which the rate of evolution of our trait is assumed to be under the influence of a second, unobserved, discrete character trait. This class of method traces its history to the covarion model of Tuffley and Steel (1997), but was adapted to phylogenetic comparative analysis as the ‘hidden-rates-model’ (HRM) by Beaulieu et al. (2013). The HRM differs from what we have proposed herein in that under the HRM we allow our data to tell us where character evolution may have changed in rate, rather than hypothesizing the location of one or more rate shifts a priori.

Finally, a completely separate class of model has recently been proposed for the phylogenetic comparative analysis of discrete character evolution that is based on the threshold model from quantitative genetics originally developed by Wright (1934; Felsenstein 2005, 2012). According to the threshold model, our observed discrete character is simply a manifestation of an underlying, unobserved continuous trait and one or more thresholds. Under this model, whenever the unobserved continuous trait crosses a threshold, the discrete character changes state. The threshold model can also create heterogeneity in the evolutionary process through time and among lineages. For instance, lineages near the threshold may change state many times, whereas clades far from any threshold will tend not to change at all (Revell 2014).

The method of this study should not be viewed as a competitor or replacement for any of the aforementioned classes of approach. To the contrary, we view it simply as one of several complementary tools whose suitability for use will depend on our data and, most importantly, on the specifics of our biological question and hypotheses.

### Notes on the empirical case studies

In addition to presenting this method and exploring its statistical attributes via simulation, we also applied the method to a pair of case studies. In each, we hypothesized that the rate of evolution for a discrete character (dominant dewlap color and number of caudal vertebrae) might differ between mainland and Caribbean *Anolis* lizard faunas. Transitions between the Central and South American mainland and the Caribbean have occurred so few times that they can be reasonably unambiguously reconstructed on the tree of anoles (Figure 3). We hypothesized a priori that dominant dewlap color might evolve at different rates in Caribbean and mainland lineages due to average (and opposite) differences in diversity and abundance in these island versus mainland environments. For example, higher densities of conspecifics might favor stronger and more frequent selection acting on dewlap coloration to compete for mates. On the other hand, higher species richness could favor the evolution of greater dewlap diversity to reduce costly heterospecific mating attempts. We also hypothesized that caudal vertebra number might evolve more rapidly in Caribbean lizards under the assumption that Caribbean anoles tend to exhibit more specialized arboreal niche use, and thus that the tail might be under stronger divergent selection on the islands due to the different locomotor demands imposed by highly specialized arboreal locomotion.

We found relatively little evidence in support of either hypothesis; however, from the perspective of methodology, perhaps our non-finding should be as much encouraging as it is disappointing. Over the past few years it has become a relatively popular endeavor to identify circumstances in which phylogenetic comparative methods can lead us spuriously astray due to null-model inadequacy (Maddison and Fitzjohn 2015; Rabosky and Goldberg 2015; O’Meara and Beaulieu 2016). Maybe we should feel encouraged the fact that not all simple null hypotheses will be rejected in favor of more complicated alternatives, if these alternatives are not, in fact, supported by our data. On the other hand, it would be incorrect to claim that our results show that no interesting differences exist in tail or dewlap evolution between mainland and Caribbean anoles. To the contrary, our analysis has merely demonstrated that a pair of relatively simple a priori hypotheses for rate variation between continental and island anoles are not supported when confronted with data.

## Conclusion

Here we have presented a relatively simple extension of the familiar M*k* model of Lewis (2001), but in which the rate or process of discrete trait evolution is permitted to differ among branches, clades, or time periods that have been hypothesized a priori by the user. We show via simulation that the method has reasonable statistical attributes – such as type I error not significantly higher than its nominal value and median parameter estimates across simulations that are very close to their generating values. Finally, we apply the method to two different case studies comparing the rate of discrete character evolution between Caribbean and mainland *Anolis* lizard clades.

## Acknowledgements

The authors would like to thank A. Harrison, R. Castañeda, R. Glor, A. Herrel, S. Poe, and Y. Stuart for their help compiling the dewlap color data used in this study; K. Winchell for providing the image used in Figure 4; and the United States National Science Foundation (DEB 1350474 and ABI 1759940) for supporting portions of this research.

---

**Table S1.**
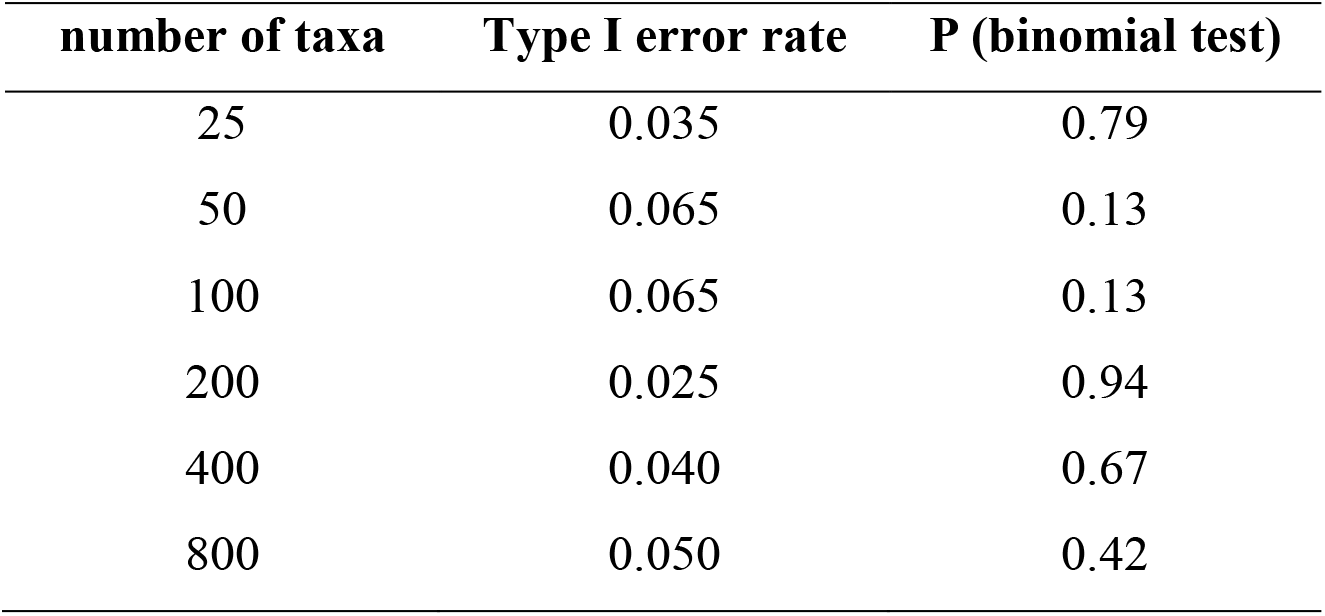
Results from an analysis of type I error. The P-values are based on a one-tailed binomial test against a type I error rate of 0.05.

**Table S2.**
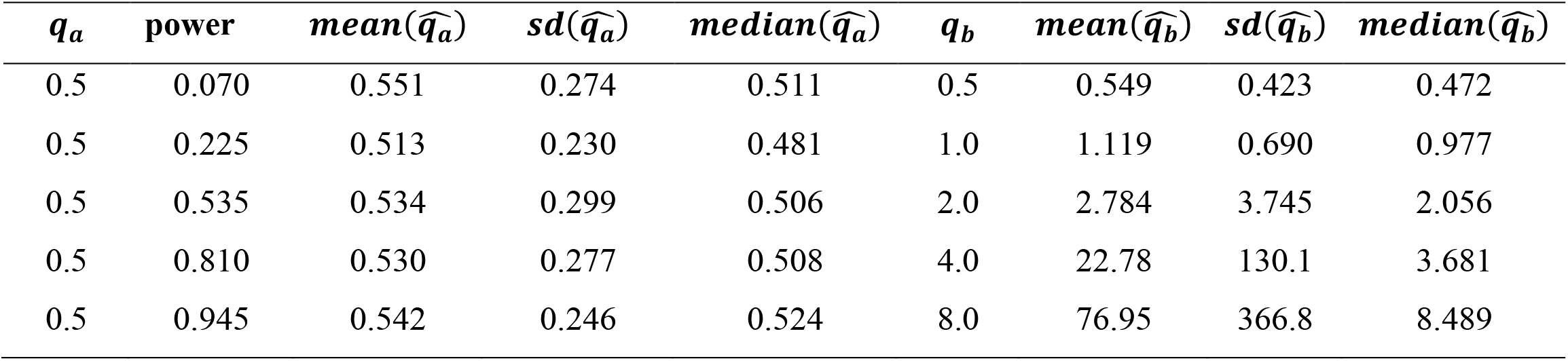
Results from an analysis of parameter estimation and power. Shown are the frequency with which the null hypothesis of equal rates was rejected (power), as well as the mean, standard deviation, and median values of the estimated parameters 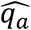 and 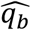.

| $q_a$ | power | $mean(\widehat{q}_a)$ | $sd(\widehat{q}_a)$ | $median(\widehat{q}_a)$ | $q_b$ | $mean(\widehat{q}_b)$ | $sd(\widehat{q}_b)$ | $median(\widehat{q}_b)$ |
| --- | --- | --- | --- | --- | --- | --- | --- | --- |
| 0.5 | 0.070 | 0.551 | 0.274 | 0.511 | 0.5 | 0.549 | 0.423 | 0.472 |
| 0.5 | 0.225 | 0.513 | 0.230 | 0.481 | 1.0 | 1.119 | 0.690 | 0.977 |
| 0.5 | 0.535 | 0.534 | 0.299 | 0.506 | 2.0 | 2.784 | 3.745 | 2.056 |
| 0.5 | 0.810 | 0.530 | 0.277 | 0.508 | 4.0 | 22.78 | 130.1 | 3.681 |
| 0.5 | 0.945 | 0.542 | 0.246 | 0.524 | 8.0 | 76.95 | 366.8 | 8.489 |

**Figure S1.**
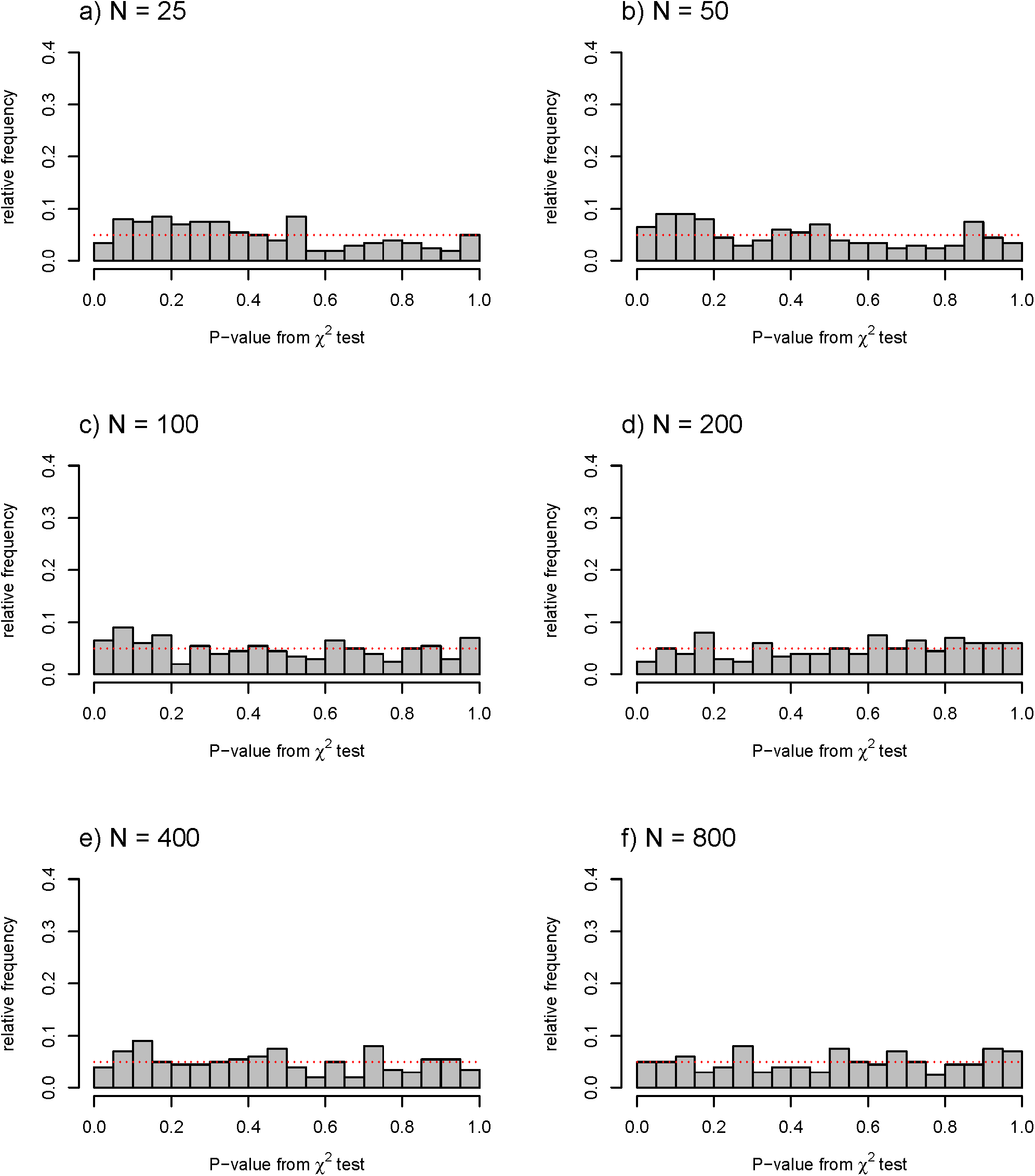
Distribution of P-values for the statistical test for data simulated under the null hypothesis. The expected distribution (uniform on the interval 0,1) is given by the horizontal dotted line in each panel.

**Figure S2.**
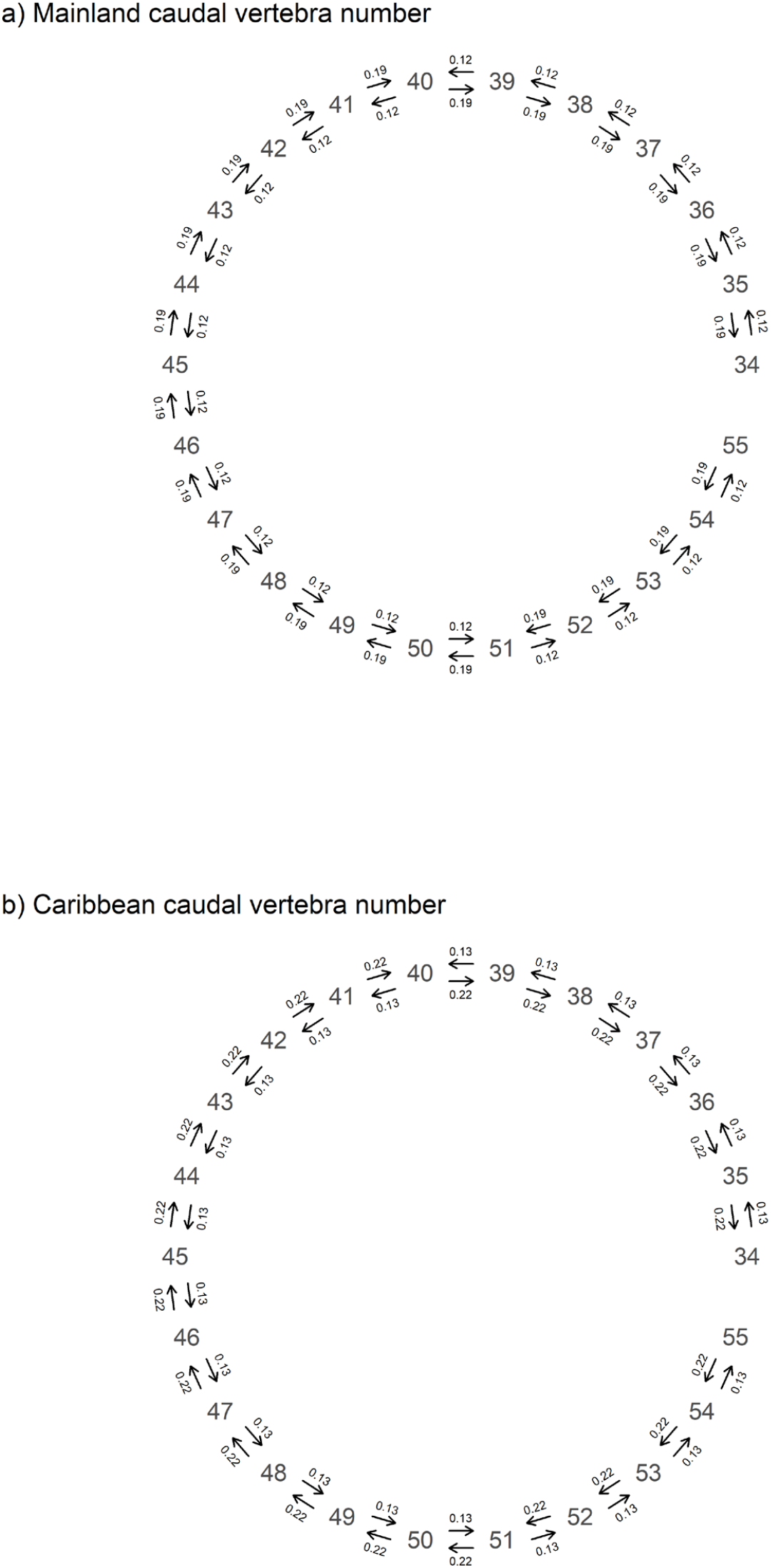
Fitted multi-rate asymmetric ordered transition model for the evolution of caudal vertebra number in mainland (panel a) and Caribbean (panel b) anole lineages.

